# PD-1 signaling uncovers a pathogenic subset of T cells in inflammatory arthritis

**DOI:** 10.1101/2023.11.16.566893

**Authors:** Johanna Straube, Shoiab Bukhari, Shalom Lerrer, Robert Winchester, Brian Henick, Matthew Dragovich, Adam Mor

**Author notes:** Equal contribution (Johanna Straube, Shoiab Bukhari). Correspondence should be addressed to: Adam Mor MD PhD 650 W 168 St. BB-1701F, Columbia University Medical Center New York, NY 10032, USA.

## Abstract

**Background:** PD-1 is an immune checkpoint on T cells and interventions to block this receptor result in T cell activation and enhanced immune response to tumors. Paired to that, and despite a decade of research, approaches to treat autoimmunity with PD-1 agonists still need to be more successful. To resolve this, new methods must be developed to augment PD-1 function beyond engaging the receptor.

**Methods:** We conducted a flow cytometry analysis of T cells isolated from the peripheral blood and synovial fluid of patients with rheumatoid arthritis. In addition, we performed a genome-wide CRISPR/Cas9 screen to identify genes associated with PD-1 signaling. We further analyzed genes involved in PD-1 signaling using publicly available bulk and single-cell RNA sequencing datasets.

**Results:** Our screen confirmed known regulators in proximal PD-1 signaling and, importantly, found an additional 1,112 unique genes related to PD-1 ability to inhibit T cell functions. These genes were strongly associated with the response of cancer patients to PD-1 blockades and with high tumor immune dysfunction and exclusion scores, confirming their role downstream of PD-1. Functional annotation revealed that more significant genes uncovered were those associated with known immune regulation processes. Remarkably, these genes were considerably downregulated in T cells isolated from patients with inflammatory arthritis, supporting their overall inhibitory functions. A study of rheumatoid arthritis single-cell RNA sequencing data demonstrated that five genes, KLRG1, CRTAM, SLAMF7, PTPN2, and KLRD1, were downregulated in activated and effector T cells isolated from synovial fluids. Back-gating these genes to canonical cytotoxic T cell signatures revealed PD-1^+^ HLA-DR^HIGH^ KLRG^LOW^ T cells as a novel inflammatory subset of T cells.

**Conclusion:** We concluded that PD-1^+^ HLA-DR^HIGH^ KLRG^LOW^ T cells are a potential target for future PD-1 agonists to treat inflammatory diseases. Our study uncovers new genes associated with PD-1 downstream functions and, therefore, provides a comprehensive resource for additional studies that are much needed to characterize the role of PD-1 in the synovial subset of T cells.

## Introduction

Ten years ago, cancer immunotherapy utilizing checkpoint inhibitors was recognized as the breakthrough of the year for its role in the paradigm shift of cancer therapies [1]. Programmed cell death protein-1 (PD-1), a receptor expressed on the T cell’s surface, antagonizes their activation [2]. PD-1 engages with two ligands, PD-L1 and PD-L2, presented in hematopoietic, non-hematopoietic, and antigen-presenting cells [3]. These interactions down-regulate the immune system and promote self-tolerance by suppressing T cell pro-inflammatory activity [4]. The emergence of checkpoint inhibitors has significantly improved the outcome of patients with advanced melanoma, non-small cell lung cancer, head and neck cancer, renal cancer, and hepatocellular carcinoma, among others [5]. As immunotherapy stimulates the body’s natural defense against tumors, it also causes immune-related adverse events resulting from misdirected immune-system stimulation in approximately 50% of patients treated with PD-1 or PD-L1 inhibitors [6]. These adverse events can be organ-specific, such as colitis, hepatitis, pneumonitis, and hypothyroidism, or nonspecific, such as fatigue, diarrhea, and rash related to immune activation [7, 8]. In addition, musculoskeletal problems have also been described in patients treated with anti-PD-1 antibodies [8]. However, their mechanisms are not entirely understood yet.

Rheumatoid arthritis (RA) is a common chronic autoimmune disease attacking synovial tissue, leading to significant morbidity and mortality [9]. With a prevalence of 1% worldwide, the etiology of RA remains poorly understood [10]. Due to progressive joint injury, individuals experience pain, limited mobility, and disability [9]. Furthermore, if left untreated, the inflammatory process can contribute to an increased risk of atherosclerosis and reduced life expectancy [11]. Despite the advice of biological therapies, many patients do not respond, and new treatments are needed. A novel approach utilizing the PD-1 pathway to treat autoimmunity envelops PD-1 agonists to inhibit T cell function. Several products have been developed already. However, only limited success has been reported in clinical trials [12].

This study identified genes potentially regulating PD-1 downstream T cell inhibition through a genome-wide CRISPR screen. Our studies have reported that genetic variations in KLRG1, CRTAM, SLAMF7, PTPN2, and KLRD1 could be linked to altered PD-1-mediated cellular immune responses and may lead to autoimmune disorders. Though much is still unknown and additional studies are needed to characterize the mechanical roles better, our findings reveal potential targets for future PD-1 agonists to treat inflammatory diseases.

## Methods

### Cell isolation, culture, stimulation

Jurkat T cells were obtained from the ATCC and maintained in RPMI medium supplemented with 10% FBS and 1% penicillin and streptomycin. Peripheral blood was acquired from the New York Blood Center. Total CD3^+^ T cells were isolated by density gradient centrifugation (Lymphoprep) and adverse selection using the RosetteSep human T cell enrichment cocktail (Stemcell). Primary T cells were directly employed in stimulation assays or maintained in culture. T cell cultures were maintained in complete RPMI, containing 10% FCS, MEM nonessential amino acids, 1mM sodium pyruvate, 100 IU/ml of penicillin, 100 μg/ml streptomycin, and GlutaMAX-I. For stimulation, Dynabeads M270-Epoxy (Thermo) was covalently conjugated with combinations of mouse anti-human CD3 antibody (clone UCHT1, BioLegend), mouse anti-human CD28 antibody (BioLegend), recombinant human PD-L1 human IgG_1_ Fc chimera protein (R&D Systems), or mouse IgG_1_ isotype antibodies (R&D Systems) following the manufacturer’s recommendations. All Jurkat and primary T cell stimulations were performed with beads at a 1:5 cell-to-bead ratio.

### Stable knockdown Jurkat T cells

Protein expression was stably knocked down in Jurkat T cells by RNA interference using Mission shRNA plasmids (Sigma-Aldrich). Lentiviral particles were generated by transfecting HEK293T cells with pMD2G, psPAX2, and the shRNA plasmid using SuperFect (Qiagen). T cells were transduced by centrifugation and selected with puromycin.

### RT-PCR analysis

Total RNA was extracted using the RNeasy Plus Mini Kit (Qiagen). RNA (500 ng) was used for cDNA synthesis using SuperScript II First-Strand Synthesis (Invitrogen). Human kinases and HPRT Taqman Primer/Probes were utilized for all Taqman Gene Expression Assays with the Taqman Universal PCR Master Mix (Applied Biosystems). Quantitative gene expression analyses were performed with Applied Biosystems 7300 Real-Time PCR. Gene expression was analyzed by the ΔΔCt method.

### Cytokine secretion

IL-2 concentrations in the supernatant were measured by enzyme-linked immunosorbent assay (ELISA) from BioLegend. For the CRISPR screen, IL-2 cytokine secretion assay (MACS) was used to label the cells and detect cytokine release.

### Western blotting

Following stimulation with CD3 beads, cells were placed on ice, resuspended ice-cold PBS, and centrifuged for 5 minutes at 400g and 4°C. The cell pellets were resuspended in cold RIPA lysis buffer containing one mM sodium orthovanadate and complete Mini, EDTA-free protease inhibitors (Roche). The cells were placed on a rotator, and lysis was carried out at 4°C for 30 minutes. The lysates were centrifuged for 10 minutes at 12,000 g and 4°C. The lysate was resuspended in reducing Laemmli buffer, boiled at 95°C for 10 minutes, and run on SDS-PAGE. Following protein transfer for 30 minutes at 25 V, the nitrocellulose membrane was blocked with 5% bovine serum albumin (BSA) in PBS containing 0.05% Tween-20 (PBST) and blotted overnight with primary antibody prepared in PBST containing 2% BSA. The membrane was developed with IRDye secondary fluorescent antibody and acquired on the Odyssey CLx Imaging system.

### Cell proliferation assay

Jurkat T cells were activated with soluble anti-CD3 or anti-CD3 + anti-CD28 cultured for 72 hours. The number of cells was assessed after 24, 48, and 72 hours by automated counting (Invitrogen Countess II) in the presence of trypan blue.

### Patient recruitment

We collected whole blood and synovial fluid from adult patients from the Columbia University rheumatoid arthritis cohort according to the approvals granted by the Columbia University Institutional Review Board.

The whole blood of healthy donors was obtained from the New York Blood Center. The Institutional Review Board at Columbia University Medical Center approved the study, and all donors provided informed consent (AAAB3287). PBMC from RA patients and healthy controls were isolated from peripheral blood using ficoll gradient centrifugation. Red blood cells were lysed by resuspending cell pellets in ACK Lysis Buffer (Gibco) for 2 minutes, followed by washing with cold PBS. PBMC were stored at −80°C until analysis.

### Flow cytometry analysis

Cells were collected and stained for protein expression analysis with the following antibodies for surface protein expression: CD4, CD8, PD-1, LAG3, TIM-1, TIGIT, ICOS, and KLRG1. Dead cells were excluded from the analysis by using Ghost Dye UV450. A complete list of antibodies used for the flow cytometry is provided as an additional file (Antibody panel). Doublets and double-positive CD4/CD8 cells were removed through sequential gating. Flow cytometry acquisition was made using the BD LSRII with BD FACSDiva. Data were analyzed by FlowJo 10.1r7 and GraphPad Prism 9.

### Library preparation

GeCKO V2 (A and B from Thermo Fisher) library consists of specific sgRNA sequences for gene knock-out in the human genome was used. The library contains six sgRNAs per gene (targeting 19,050 genes) for adequate representation and 1,864 control sgRNAs designed not to target the genome. The library was provided in one vector (lentiCRISPRv2) format. These vectors enable lentiviral delivery of both Cas9 and sgRNA for targeted gene knockout. To generate lentivirus, lentiCRISPR (with sgRNA cloned) was co-transfected into HEK293T cells with the packaging plasmids pVSVg (AddGene 8454) and psPAX2 (AddGene 12260). As a positive control for viral production, we used CMV-EGFP lentiviral transfer plasmid (AddGene 19319). The library contains 122,411 (65,383 in Library A, 58,028 in Library B) unique sgRNAs. The library was diluted to 50 ng/uL in water and electroporated to competent (NEB DH5a cells) cells with an efficiency of ≥10^9^ cfu/ug. A 10 cm petri dish (ampicillin) was used for 40,000-fold dilution of the entire transformation to estimate transformation efficiency and to ensure that full library representation is preserved. Plates were grown for 14 hours at 32°C. Calculate transformation efficiency by counting the number of colonies on the dilution plate. Next, we harvested the colonies and performed Maxi-prep for downstream virus production and future amplification.

### Pooled genome-wide CRISPR screen

Jurkat T cells were cultured in RPMI with 10% FBS. Cells were infected with the pooled lentiviral library at an MOI of 0.3 to ensure that only one gene was targeted for Cas9-mediated editing in each cell. To fully represent the library sgRNA sequences in the transduced cell population, the library coverage at transduction was determined to be ∼100 transduced cells for each sgRNA. The transduced cells were selected for three days with puromycin (2 μg/mL). Following puromycin selection, the cell populations were maintained at a low concentration (1 million cells/mL). For the genome-wide screen, 500 million transduced Jurkat T cells were stimulated with anti-CD3/CD28, with and without recombinant PD-L1, as described above. IL-2^HIGH^ and IL-2^LOW^ cell populations were sorted using MACS according to the manufacturer’s instructions. Cells were resuspended in 20 mL P1 buffer (Qiagen) with 100 μg/mL RNaseA and 0.5% SDS. After incubating at 37°C for 30 minutes, the lysate was heated at 55°C for 30 minutes in the presence of Proteinase K (100 μg/mL). After digestion, samples were passed through a needle multiple times. Next, 20 mL Phenol: Chloroform: Isoamyl Alcohol (Invitrogen #15593–031) was added into homogenized samples. After mixing, the samples were transferred into 50 mL MaXtract tubes (Qiagen) and centrifuged at 1,500 × *g* for 5 minutes at room temperature. The aqueous phase was transferred into ultracentrifuge tubes and thoroughly mixed with 2 mL 3 M sodium acetate plus 16 mL isopropanol at room temperature before centrifugation at 15,000 × *g* for 15 minutes. The gDNA pellets were carefully washed with 10 mL of 70% ethanol and dried at 37°C. Dry pellets were resuspended in water, and gDNA concentration was adjusted to 1 μg/uL. PCR reactions were prepared to allow amplification of the total harvested gDNA from a 1,000-cell coverage for each sample using the following primers: GGCTTGGATTTCTATAACTTCGTATAGCA; CGGGGACTGTGGGCGATGTG; AATGATACGGCGACCACCGA; GATCCACAAAAGGAAACTCACCCTAAC; CAAGCAGAAGACGGCATACGAGAT. The resulting PCR product (344 bp) was extracted from a 1% agarose gel. Gel-extracted bands were submitted for sequencing on an Illumina HiSEq.

### Dataset selection, exploration, and visualization

The criteria for selecting the downstream analysis datasets were T-cell-focused studies, inflammatory arthritis, or immunotherapy studies. The siRNA sequencing datasets consisting of cells from patients with cutaneous melanoma (SKCM) GSE166181 (48 patients), basal cell carcinoma (BCC) GSE123813 (11 patients), and squamous cell carcinoma (SCC) GSE123813 (3 patients) were assessed and analyzed by TISCH2 database (http://tisch.comp-genomics.org). The siRNA sequencing datasets consisting of patients with metastatic melanoma (GSE120575; 32 patients), metastatic renal cell carcinoma SCP1288 (8 patients), melanoma GSE164237 (4 patients), and arthritis MTAB9492 (5 patients) were investigated, and visualized using Bioturing (https://bioturing.com). The Seurat package confirmed the quality of scRNA datasets and reannotated cell states. Unsupervised clustering of cells from a given dataset was performed using the standard pipeline. The first 25 principal components (PCs) were used for Louvain graph-based clustering implemented in the Seurat package. Uniform manifold approximation and projection (UMAP) was performed on the same PCs with 100 nearest neighbors for visualization in two dimensions the following bulk RNA dataset of the patients with RA: GSE65010 (6 controls and six patients), GSE56649 (9 controls and 13 patients), GSE38351 (31 controls and 12 patients), GSE57383 (19 controls and nine patients), GSE110169 (77 controls and 84 patients), GSE90081 (12 controls and 12 patients). These datasets were downloaded from GEO NCBI (https://www.ncbi.nlm.nih.gov/geo). All the datasets were analyzed separately and as integrated and normalized (z-scored) where mentioned. The gene expression heatmaps were generated using Morpheus (https://morpheusdata.com) or Bioturing (https://bioturing.com). A based method was implemented to analyze and generate a differential expression matrix via empirical Bayes moderated t-statistics and associated p-values, as represented by the volcano plots. ShinyGO (http://bioinformatics.sdstate.edu) developed the gene ontology enrichment analysis). Beeswarm plot was used to show immune-related enriched genes (https://www.indiegogo.com). The list of the transcription factors related to immune function was acquired from the MSigDB dataset (https://www.gsea-msigdb.org/gsea). Some plots were generated by Prism (GraphPad) and Talk2data-Bioturing (https://talk2data.bioturing.com/login).

### Statistical analysis

Statistical significance was determined using ordinary 1-way ANOVA, Tukey’s multiple comparisons test with a single pooled variance, or post-hoc paired test applying Šidák’s multiple comparisons test wherever indicated. Statistical analyses were performed using Prism 9 (GraphPad Software). Significance was set at p = 0.05.

## Results

### PD-1 levels in synovial T cells are higher than in peripheral blood T cells

Although PD-1 is expressed on activated T cells and some subsets of memory T cells, PD-1 agonist interventions have not worked as expected in treating autoimmune diseases like lupus and rheumatoid arthritis (RA). To answer whether this is due to different expression levels of PD-1 in the peripheral blood (PB) versus the synovial fluid (SF) T cells in RA patients, we collected synovial fluids and blood from RA patients and peripheral blood from healthy controls (HC). We measured PD-1 expression on different subsets of T cells by flow cytometry. As shown (Fig. 1A-D), PD-1 levels were higher on synovial CD4 and CD8 effector memory T cells than peripheral blood T cells isolated from the same RA patients and HC. Quantifying the percentages of PD-1 positive CD4 and CD8 T cells revealed a higher proportion of RA synovial fluid T cells than those isolated from RA or HC peripheral blood (Fig. 1Ei and 1Fi). T cell subset analysis revealed that most PD-1 positive T cells were effectors and central memory cells (Fig. 1Eii and 1Fii). Moreover, these T cells tested positive for the checkpoints LAG3, ICOS, and TIGIT (Fig. 1Eiii and 1Fiii). This data suggests that PD-1 is highly expressed in RA T cells, occupying mainly the synovial fluid compartment.

**Figure 1.**
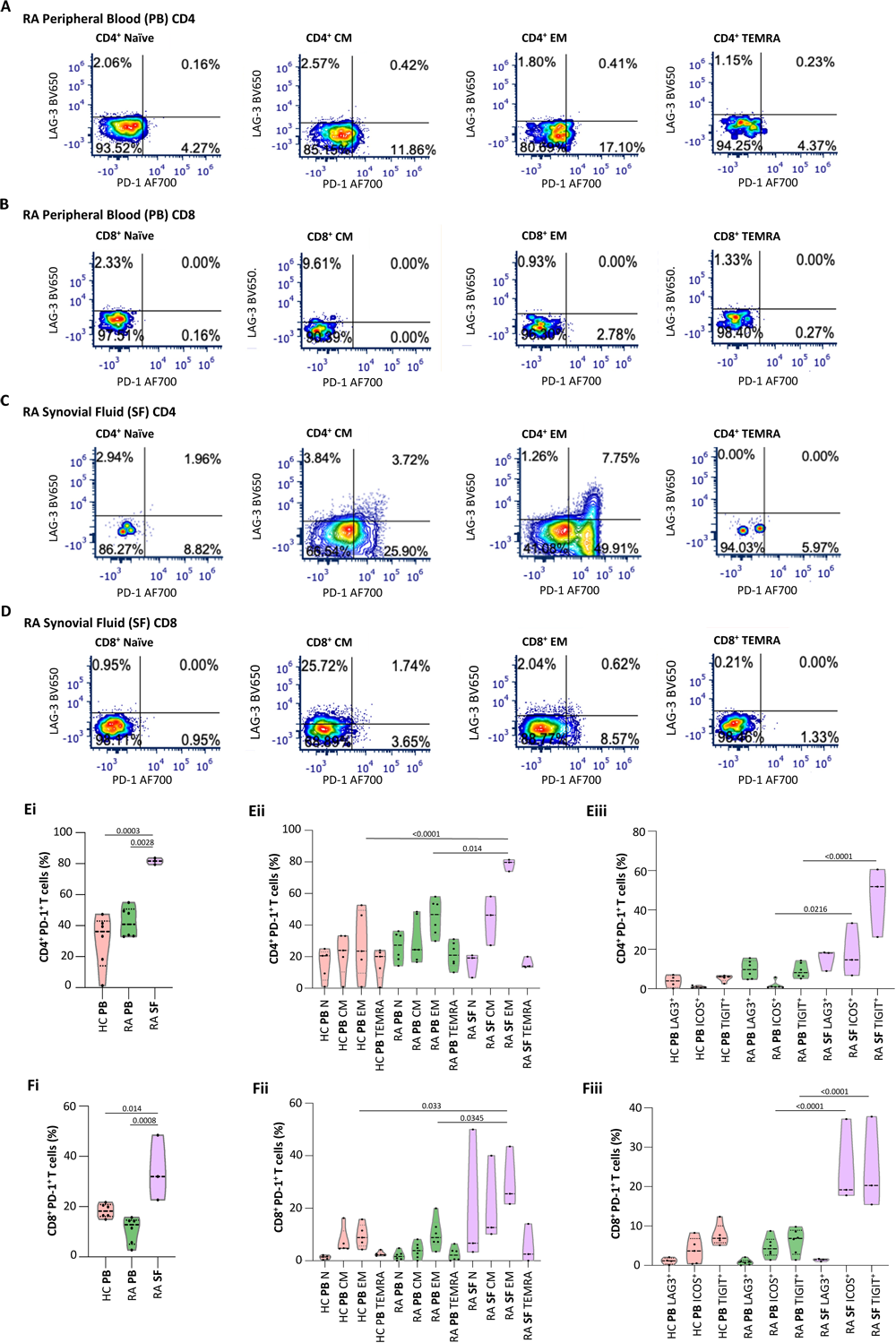
PD-1 levels in synovial T cells are higher than in peripheral blood T cells. Flow cytometry was performed on T cells isolated from synovial fluid (SF) and matched peripheral blood (PB) T cells from patients with Rheumatoid arthritis (RA) (A-D). Flow cytometry analysis of the expression levels of PD-1 and LAG-3 was performed on naïve (N), central memory (CM), effector memory (EM), terminally differentiated memory cells (TEMR) CD4, and CD8 T cells (A-D). Violin plots quantification of the percentages of CD4^+^PD-1^+^ cells in healthy controls (HC) PB, RA PB, and RA SF out of all CD4^+^ cells (Ei). Violin plots quantification of the percentages of CD4^+^PD-1^+^ cells in HC PB, RA PB, and RA SF out of all CD4^+^ cells within N, CM, EM, and TEMRA subsets (Eii). Violin plots quantification of the percentages of CD4^+^PD-1^+^ cells expressing LAG3, ICOS, or TIGIT in HC PB, RA PB, and RA SF out of all CD4^+^PD-1^+^ cells (Eiii). Violin plots quantification of the percentages of CD8^+^PD-1^+^ cells in healthy controls HC PB, RA PB, and RA SF out of all CD8^+^ cells (Fi). Violin plots quantification of the percentages of CD8^+^PD-1^+^ cells in HC PB, RA PB, and RA SF out of all CD8^+^ cells within N, CM, EM, and TEMRA subsets (Fii). Violin plots quantification of the percentages of CD8^+^PD-1^+^ cells expressing LAG3, ICOS, or TIGIT in HC PB, RA PB, and RA SF out of all CD8^+^PD-1^+^ cells (Fiii). Statistical significance was determined using ordinary 1-way ANOVA, Tukey’s multiple comparisons test; * p<0.05, n=3-5.

### A genome-wide CRISPR screen uncovered genes associated with PD-1 inhibitory function

To better understand some of the reasons for the limited response to PD-1 agonists in RA and to uncover novel PD-1 downstream therapeutic targets, we performed a genome-wide CRISPR Cas9 screen using Jurkat T cells stimulated with anti-CD3 and anti-CD28 antibodies with and without recombinant PD-L1. Stimulation of T cells with anti-CD3 and anti-CD28 antibodies leads to increased IL-2 secretion, but engaging PD-1 with PD-L1 averts that, leading to a reduction in the rate of IL-2 secretion. T cells were infected with a lentivirus library containing 64,012 guides directed to 21,342 genes, including 1,350 nontargeting control sequences (Fig. 2A). Loss of ability of PD-1 signaling to inhibit IL-2 secretion was identified by flow cytometry, and cells with high levels of IL-2 (cells in which PD-1 failed to inhibit IL-2 secretion) and low levels of IL-2 (cells where PD-1 signaling inhibited IL-2 secretion) were sorted (Sup. Fig. 1). CRISPR guide sequences from both populations of cells were PCR amplified, and NGS sequenced. We discovered 1,080 genes that were not involved in PD-1 downstream signaling and 1,112 genes that were associated with PD-1 downstream signaling (Fig. 2B) (Sup. Tab. 1). In a secondary validation screen, we chose 22 genes generated shRNA knocked down Jurkat T cell lines and validated the lack of ability of PD-1 to inhibit IL-2 secretion using ELISA (Fig. 2C). Furthermore, we confirmed that in PTPN2 deficient T cells, additional inhibitory functions of PD-1 were abrogated (Sup. Fig. 2) and that in FUT6 knocked down cells, T cell receptor, and PD-1 signaling were impaired (Sup. Fig. 3).

**Figure 2.**
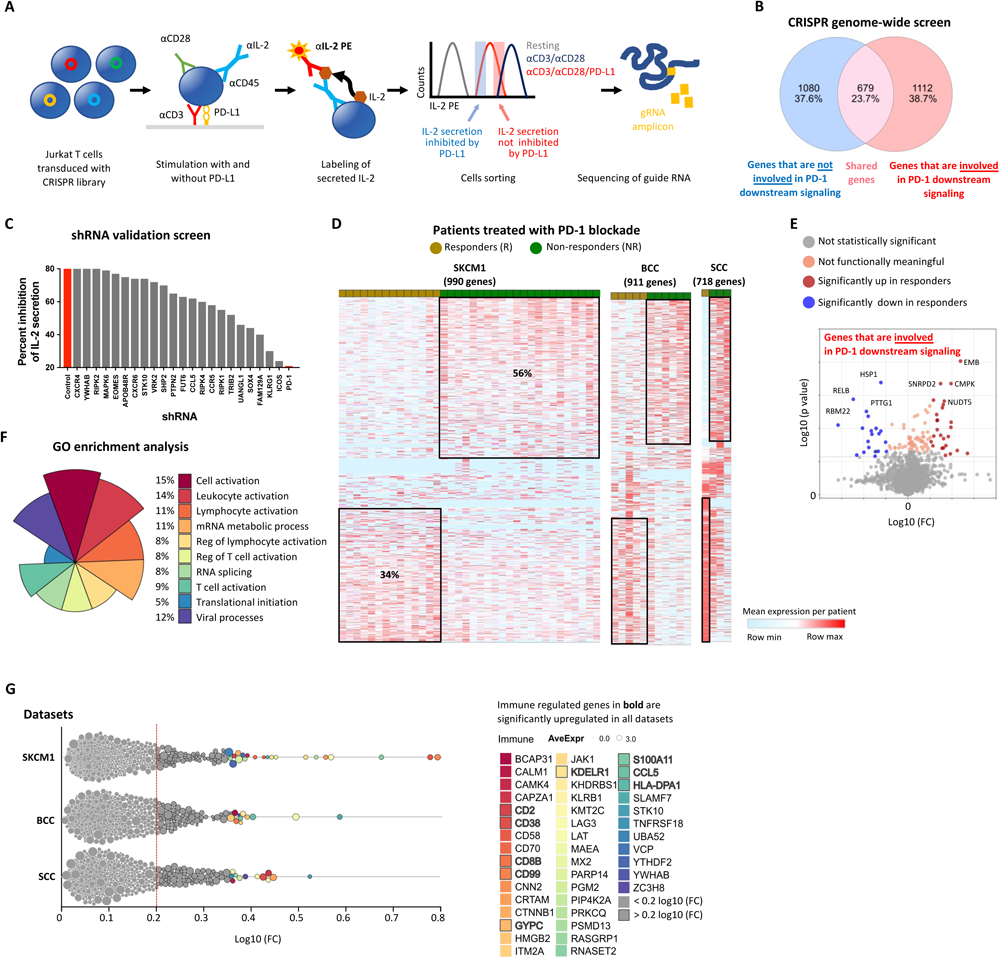
The expression of genes required for PD-1 signaling is associated with clinical response to PD-1 blockade. Diagram of the experimental pipeline from transduction of the Jurkat T cells, cell labeling, stimulation, sorting, and guide sequencing (A). Venn diagram showing the number of genes identified by sequencing in the two groups of sorted cells (B). Graph showing the levels of secreted IL-2 measured in each clone of Jurkat short hairpin knockdown T cells under the indicated treatment conditions, n=4 for each cell line; average values are shown (C). Heatmaps showing the differential expression levels of genes that we discovered in our screen to be involved in PD-1 downstream signaling, now in T cells isolated from cancer patients that were treated with anti-PD-1 antibodies in three separated clinical trials (SKCM1, BCC, SCC) (D). The patients were divided into those who responded favorably to PD-1 blockade (R) and those who failed to respond (NR). Volcano plots displaying expression levels of genes that were either up or downregulated in the cancer patients that failed to respond to PD-1 blockade (E). Differences in the expression levels were calculated by implementing Limma statistics. GO-enrichment analysis of the genes identified in the screen and involved in the PD-1 signaling pathway (F). A berswarm plot expression levels of genes discovered in the screen and as differentially expressed in the T cell from the cancer patients subjected to PD-1 blockade (G). Bolded genes are those that were shared in all three clinical trials.

### The expression of genes required for PD-1 signaling is associated with clinical response to PD-1 blockade

We discovered 1,080 genes that were not involved in PD-1 downstream signaling and 1,112 genes that were engaged in PD-1 downstream signaling (Fig. 2B). To assess the relevancy of PD-1-associated top-ranked genes, we analyzed the expression levels of these genes in the anti-PD-1 treated cancer datasets. We selected three cancer immunotherapy single-cell RNA sequencing datasets comprising 51 cancer patients and 95,347 T cells (Fig. 2D) (Sup. Fig. 4). Interestingly, most of our top-ranked PD-1-associated genes were differentially expressed in T cells isolated from cancer patients that did or did not respond favorably to PD-1 blockade. 56% of the top-ranked genes were upregulated in the non-responders (NR) compared to 34% of the top-ranked genes that were upregulated in the responders (R) (Fig. 2D). Remarkably, genes that have been reported not to be associated with PD-1 signaling or other inhibitory T cell functions, such as RELB and HSP1, were significantly lower in the NR cancer patients (Fig. 2E). GO-enrichment analysis confirmed that the genes that we identified as involved in PD-1 signaling were associated with T cell inhibitory functions (Fig. 2F) (Sup. Tab. 2). Many of the genes that were involved in PD-1 signaling, such as CD2, CD38, CD99, KDELR1, and S100A11 were found to be highly expressed in tumor-infiltrating CD8 T cells in all the three cancer datasets as shown in the beeswarm plot (Fig. 2G). Thus, this data suggests that PD-1-involved genes are associated with a tumor response to PD-1 blockade among cancer patients.

### Genes that are associated with PD-1 signaling are downregulated in inflammatory arthritis

To test the relevancy of our data regarding the signaling of PD-1 to patients with inflammatory arthritis where T cells are chronically activated, we interrogated publicly available autoimmune RNA sequencing datasets (Fig. 3A). We screened both bulk (Fig. 3B) and signal-cell RNA sequencing datasets (Sup. Fig. 5A) of T cells isolated for peripheral blood (PB) and synovial fluids (SF) of patients with active inflammatory arthritis. Interestingly, 30 of our top-ranked genes that were upregulated in the cancer NR had high tumor immune dysfunction and exclusion (TIDE) scores (Sup. Fig. 5B) and were downregulated in the T cell compartment in patients with active arthritis (Fig. 3B). This suggests that alteration in PD-1 signaling is associated with T cell activation in patients with inflammatory arthritis. Moving to inflammatory arthritis single-cell RNA sequencing data, we re-annotated cell clusters based on the genes we discovered to be involved in PD-1 signaling (Fig. 2B and Fig. 3C). Next, we assessed the expression of these genes in major subsets of CD8 T cells expressing CD69 (a marker of early activation) or HLA-DRB1 (a marker of late activation) (Fig. 3D) and effector CD4 and CD8 T cells expressing PRF1 (a feature of effector cells) (Fig. 3E). Genes commonly associated with T cell activation, such as LAG3, CCL5, ITM2A, CCR5, and CD2, were highly expressed in SF, in comparison to PB T cells of arthritis patients. Interestingly, genes such as KLRG1, LAT, PTPN2, CRTAM, and SLAMF7 that are known to regulate T cell functions were downregulated in activated CD8 T cells in SF (Fig. 3D). Next, we determined if this also holds in effector CD4 and CD8 T cells. While most T cell activation-associated markers were highly expressed in the PRF1-expressing CD4 and CD8 T cells (Fig. 3E), only effector CD8 T cells expressed lower levels of SLAMF7 and KLRG1 in the SF compartment (Fig. 3F). To gain better insight into the regulation of the PD-1-associated genes and to dissect the transcriptional regulation of T cells, we analyzed the expression levels of exhaustion markers and associated immune function-associated transcription factors (TF) (Supp. Tab. 3) in different PB and SF T cell subsets (Sup. Fig. 6). We identified variations in the expression of many TF, such as ZNF202 and SP1, among other distinct T cell subsets. We observed differences in TF occupancy, with EGR1 and SP2 being among the TF mapped using the 58 top-ranked genes associated with PD-1 signaling (Sup. Fig. 4C). Altogether, this data suggests that the alteration in PD-1 signaling is significantly associated with activating specific T cell subsets in arthritis patients.

**Figure 3.**
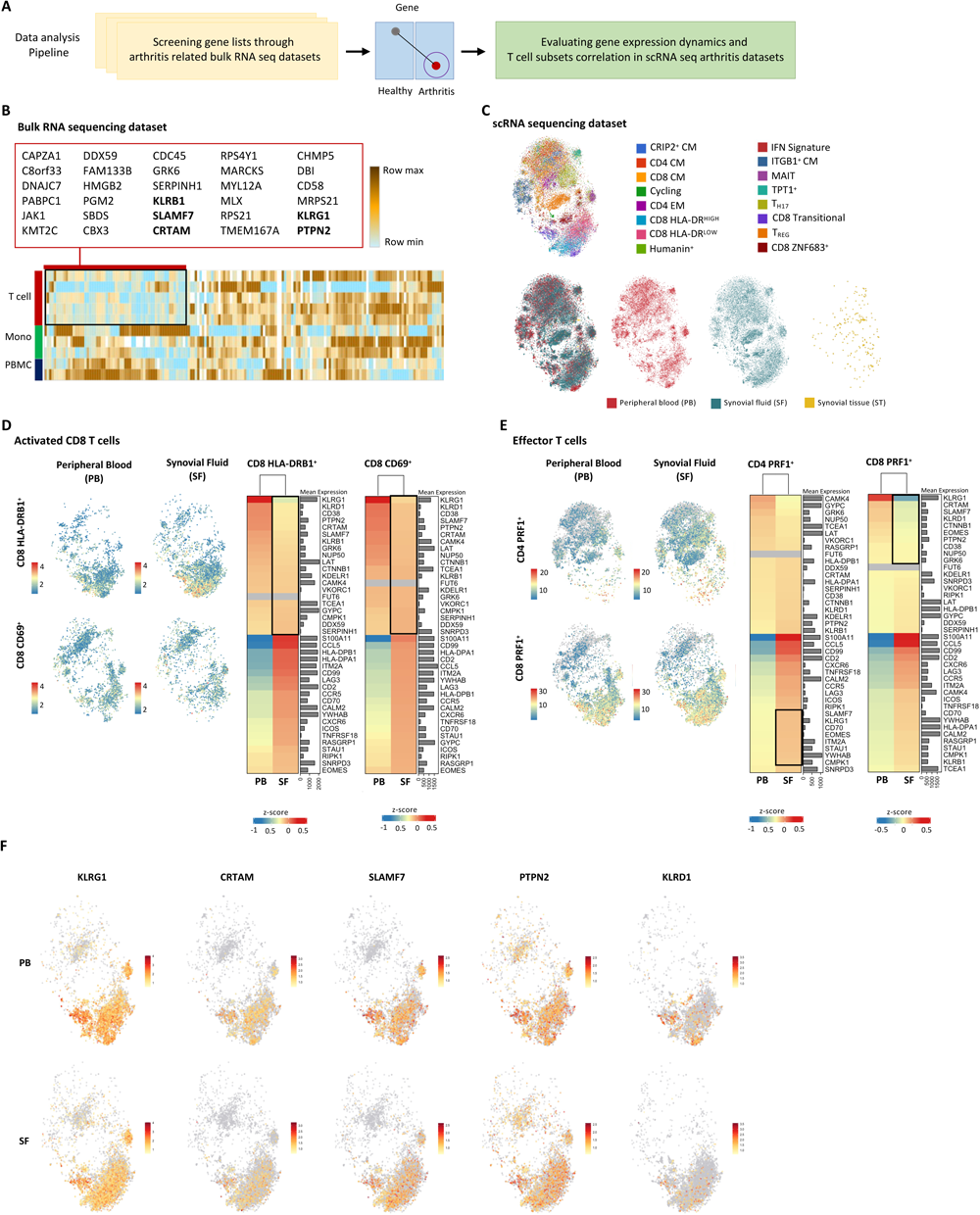
Genes that are associated with PD-1 signaling are downregulated in inflammatory arthritis. Schematic of data analysis pipeline showing the transition from the investigation of bulk RNA sequencing data to single-cell RNA sequencing data of patients with inflammatory arthritis (A). Heatmap showing RNA expression levels of top genes of bulk RNA sequencing datasets of RA patients sorted based on highest TIDE scoring (B). Highlighted in the box are downregulated genes, specifically in the T cells of RA patients. Genes in bold letters were also significantly downregulated in subsequent analysis. UMPAP plot showing clusters reannotation of single-cell RNA sequencing data of RA patients, prioritizing the genes emphasized in Figure B (C). Cells were clustered based on the T cell subset and by anatomical origin: PB, SF, or synovial tissue (ST). UMAP plots comparing the distribution of two clusters of activated CD8 T cells, CD8 HLA-DRB1^+^ and CD8 CD69^+^, isolated from either PB or SF (D). The expression of genes significantly differentially expressed in the two subsets of activated CD8 T cells is PB compared to SF. UMAP plots comparing the distribution of effector CD4 T cells (CD4 PEF1^+^) to effector CD T cells (CD4 PEF1^+^) (E). Expression of genes significantly differentially expressed between the CD4 PEF1+ and the CD8 PEF1+ effector subset in the PB and SF compartments. UMAP plots showing cell subsets expressing high levels of KLTG1, CRTAM, SLAMF7, PTPN2, and KLRD1 (F).

### Identification of a population of pathogenic PD-1^HIGH^ KLRG1^LOW^ T cells in synovial fluid

Focusing on the expression of checkpoint receptors and the top genes that were associated with PD-1 signaling highlighted the inverse correlation between these genes in both the PB and the SF, specifically in HLA-DR^HIGH^ CD8-activated T cells (Fig. 4A). Volcano plot analysis of all the genes that were expressed in this cell subset revealed that many genes linked to T cell activation were upregulated in the SF compartment, including genes such as GZMB, LMNA, and DUSP4 (Fig. 4B). Moreover, comparison between PB and SF revealed that these genes were highly expressed in the latter populations (Fig. 4C). Back gating on cells that express similar levels of KLRG1 in both compartments (Fig. 4D) detected additional genes that were only upregulated on the SF T cells (Fig. 4E), including LMNA, GZMB, and JUN (Fig. 4F). Next, we compared the expression levels of genes between cytotoxic CD8 KLRG1^LOW^ and CD8 KLRG1^HIGH^ T cells in the SF (Fig. 4G). We discovered that TIMP1 and ZNF683 were unique to the cells that express low levels of KLRG1 (Fig. 4H and 4I).

**Figure 4.**
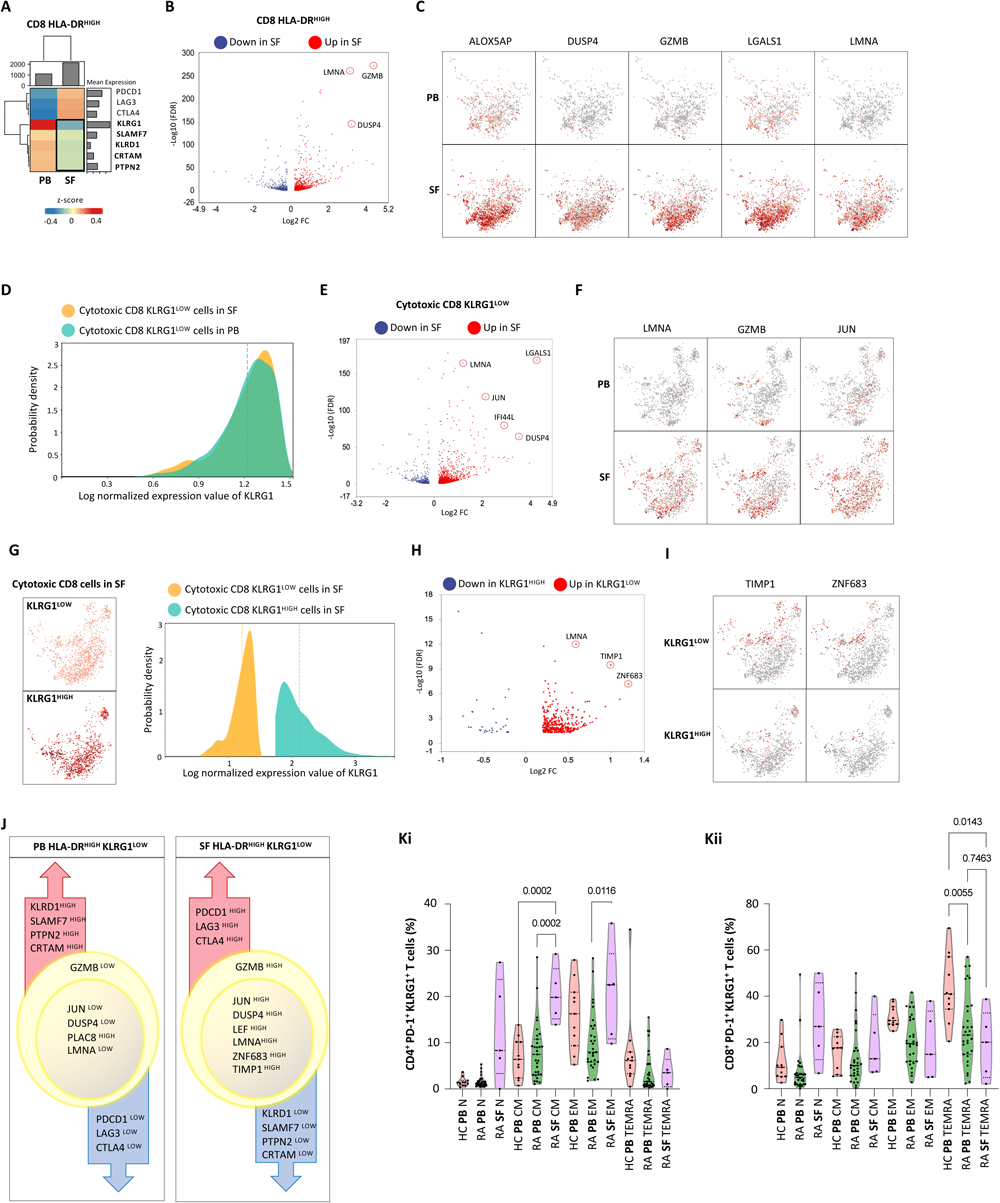
Identification of a population of pathogenic PD-1^HIGH^ KLRG1^LOW^ T cells in synovial fluid. Heatmap showing differential RNA expression levels of selected genes between PB and SF within the CD8 HLA-DR^HIGH^ T cells (A). Volcano plot showing genes that were up or downregulated in SF HLA-DR^HIGH^ cells (B). UMAP plots show the enriched population of cells expressing the genes upregulated in SF CD8 HLA-DR^HIGH^ T cells, now compared to PB CD8 HLA-DR^HIGH^ T cells (C). Density plot comparing the distribution of the expression levels of KLRG1 in cytotoxic CD8 KLRG1^LOW^ cells from SF and PB (D). Volcano plot displaying up-or downregulated genes in SF cytotoxic CD8 KLRG1^LOW^ cells (E). UMAP plots show the enriched population of cells expressing the genes upregulated in SF cytotoxic CD8 KLRG1^LOW^ SF compared to PB (F). UMAP plot and density plot showing the populations and the spectrum of expression of KLRG1 in SF cytotoxic CD8 KLRG1^LOW^ and SF cytotoxic CD8 KLRG1^HIGH^ cells (G). Volcano plot displaying genes downregulated in cytotoxic CD8 KLRG1^HIGH^ cells and upregulated in cytotoxic CD8 KLRG1^LOW^ in SF (H). UMAP plots show the enriched population of cells expressing TIMP1 and ZNF683 in cytotoxic CD8 KLRG1^LOW and^ cytotoxic CD8 KLRG1^HIGH^ SF cells (I). Diagram showing the distinctive genes up or downregulated in HLA-DR^HIGH^ KLRG1^LOW^ T cells from PB and SF (J). Violin plots summarizing the percentages of CD4^+^PD-1^+^KLRG^+^ T cells isolated from HC PB, RA PB, and RA SF samples and gated on N, CD, EM, and TEMRA subsets (Ki). Violin plots summarizing the percentages of CD8^+^PD-1^+^KLRG^+^ T cell isolated from HC PB, RA PB, and RA SF samples and gated on N, CD, EM, and TEMRA subsets (Kii). Statistical significance was determined using ordinary 1-way ANOVA, post-hoc test applying Šidák’s multiple comparisons; * p<0.05, n=5-30.

These findings suggest that cytotoxic CD8^+^ PD-1^+^ HLA-DR^HIGH^ KLRG1^LOW^ T cells in the SF are different than the same cells from the PB (Fig. 4J) and, accordingly could mediate some elements of synovial inflammation in RA and also serve as a target for therapy in the context of the PD-1 agonism. Indeed, flow cytometry analysis of PB and SF cells from RA patients and HC revealed a low level of KLRG1 positive cells in the CD4 and the CD8 compartments, specifically among the TEMRA subset (Fig. 4Ki and 4Kii).

## Discussion

Checkpoint inhibitors have significantly improved cancer patients’ survival rates [13–16]. However, most cancer patients do not respond, and many develop side effects, such as immune-related adverse events (irAEs) [7]. On the contrary, PD-1 agonists activate the PD-1 pathway [17] and, therefore, are being investigated as a potential treatment for various autoimmune diseases [17, 18]. The exact mechanism by which PD-1 agonists suppress T cell responses has yet to be fully understood. PD-1 agonists are still in the early stages of clinical development. Peresolimab, for example, is currently in phase III trials to treat RA [19, 20]. Another drug called Rosnilimab is in phase II trials to treat RA [21]. Furthermore, ImmTAAI molecules are bispecific agonists currently in phase I trials for treating SLE. The molecules are designed to target PD-1 on T cells in specific tissues, which could help to reduce the risk of side effects [18]. Other PD-1 agonists in clinical trials for autoimmune diseases include BMS-986019, BGB-A317, and TPI-201. The results of these clinical trials are still pending, and there is no guarantee that they will successfully treat autoimmune diseases. The safety of PD-1 agonists is still being evaluated. The most common side effects of PD-1 agonists are fatigue, diarrhea, and skin reactions [22]. More severe side effects, such as the reactivation of latent infections, have also been reported [23, 24]. PD-1 agonists can hypothetically trigger secondary autoimmune responses [18].

There are a few reasons why PD-1 agonists might increase cancer risk. Activation of the PD-1 pathway can inactivate tumor-infiltrating lymphocytes, thus evading immune surveillance (25). Moreover, PD-1 agonists could increase the expression of PD-L1 on cancer cells, making these cells more resistant to T cell-mediated killing. In addition, and similar to other DMARS, PD-1 antagonists could be associated with increased cancer risk in patients with RA [26]. While the risk of cancer associated with PD-1 agonists is still being studied, it is essential to understand the pathway downstream of PD-1 in T cells better to find alternative agonist therapeutic targets.

We discovered genes that regulate PD-1 downstream signaling through a genome-wide CRISPR screen. Our screen uncovered 1,112 genes required for PD-1 to inhibit T cell functions. Remarkably, many of these genes were downregulated in T cells isolated from patients with inflammatory arthritis, supporting their overall inhibitory functions. Of these genes, KLRG1, CRTAM, SLAMF7, PTPN2, and KLRG1 were downregulated in activated effector T cells isolated from SF. Through a data mining strategy, we uncover a new subset of T cells, CD8^+^ PD-1^+^ HLA-DR^HIGH^ KLRG1^LOW^, that might be associated with PD-1 downstream functions in SF and, therefore, a target for future PD-1 agonists. Through a flow cytometry analysis, we confirmed the presence of this unique subset of T cells in the SF that we collected from RA patients, which is likely to be pathogenic.

KLRG1 (killer cell lectin-like receptor G1) is an inhibitory protein expressed on the surface of T cells and natural killer (NK) cells. In general, KLRG1^LOW^ cells are supposed to be more recently activated than KLRG1^HIGH^ cells and are also considered more responsive to new antigens. KLRG1^LOW^ cells have been implicated in several immune responses, including the response to viral infections, cancer, and autoimmune diseases [27]. We discovered that activated effector CD8 T cells present in the RA SF, not in the PB, could be further characterized by low expression levels of KLRG1. These cells also express low levels of PTPN2, SLAMF7, and CRTAM.

Interestingly, low KLRG1, PTPN2, and SLAMF7 levels have been observed in many autoimmune diseases, usually in the context of tissue-resident T cells [28]. A higher percentage of SLAMF7^+^ CD8^+^ T cells has been associated with T cell exhaustion. To gain a deeper understanding of the variation in T cell states at the transcriptional level, we conducted a comparative TF analysis between activated CD8 T cells gating on cells that express high and low levels of KLRG1. We observed that even though all the cells were collected from the same anatomical compartment (SF) and shared high levels of HLA-DRB1, PRF1, GZMA, GZMB, TBX21, and CD69, they exhibited distinct transcriptional profiles based on KLRG1 levels. We observed higher expression levels of ZNF683 and TIMP1 in KLRG1^LOW^ T cells, similar to what has been reported in other tissue-resident T cells [29].

Our study has several strengths. The signaling pathways downstream of PD-1 need to be better understood, and in this context, our study provides a preliminary resource to uncover additional signaling components. The binding of PD-L1 to PD-1 phosphorylates the tyrosine in the ITSM and ITIM domains of PD-1, recruits SHP-1 and SHP-2, and inhibits the activation of PI3K/Akt, which ultimately weakens the activation and proliferation of T cells. Besides, PD-1 can block the Ras/MEK/ERK pathway, thereby regulating cell cycle molecules and halting the proliferation of T cells. Indeed, our study uncovers multiple GEF and GAP that govern the function of Ras and other small GTPases.

Our study is not free of limitations. In our validation cohort, we could not confirm the presence of tissue-resident PD-1^+^ KLRG1^LOW^ cells in both CD4 and CD8 compartments. Also, we could not demonstrate that these cells actively mediate the course of the synovial inflammatory responses. Overall, this study uncovers a unique subset of T cells where many of the signaling proteins downstream of PD-1 are differentially expressed, positioning these cells as a target for therapy to enhance or inhibit PD-1 functions.

## Conclusions

This study sheds light on the subset of activated CD8 T cells with the RA SF. Low KLRG1, PTPN2, SLAMF7, and CRTAM expression levels characterize these T cells. Notably, our transcriptional analysis reveals that despite sharing specific key markers, such as HLA-DRB1, PRF1, GZMA, GZMB, TBX21, and CD69, with other CD8 subsets, these cells exhibit unique transcriptional profiles based on KLRG1 expression. These insights enhance our understanding of the heterogeneity within CD8 T cell populations in RA SF and provide valuable groundwork for future investigations into their functional significance in autoimmunity.

### List of abbreviations

RA: Rheumatoid arthritis
SF: Synovial fluid
PBMC: Peripheral blood mononuclear cells
KLRG1: Killer cell lectin-like receptor G1
GZMA: Granzyme A
GZMB: Granzyme B
PTPN2: Protein tyrosine phosphatase non-receptor type 2
CRTAM: Class I restricted T cell-associated molecule
SLAMF7: Signaling lymphocytic activation molecule family member 7
HLA-DRB1: Human leukocyte antigen DRB1

## Declaration

### Ethics approval and consent to participate

The Institutional Review Board at Columbia University Irving Medical Center approved the study. All the donors provided informed consent (Committee’s reference number IRB-AAAB3287).

### Consent for publication

Not applicable.

### Availability of data and materials

The datasets used and analyzed during the study are publicly available and can be retrieved by inputting the following GSE dataset identifiers from the GEO database: GSE166181, GSE123813, GSE120575, and GSE164237 (https://www.ncbi.nlm.nih.gov/gds). Dataset SCP1288 is available for analysis on the single-cell portal of the Broad Institute (https://singlecell.broadinstitute.org/single_cell/study/SCP1288/tumor-and-immune-reprogramming-during-immunotherapy-in-advanced-renal-cell-carcinoma#study-download). Dataset MTAB9492 is available at EMBL-EBI (https://www.ebi.ac.uk/biostudies/arrayexpress/studies/E-MTAB-9492) and the European Genome-Phenome Archive (https://ega-archive.org/studies/EGAS00001002104).

### Authors’ contribution

SJ carried out the cellular and biochemical studies, participated in the flow cytometry and genetic assays, and drafted the manuscript. BS conducted the data analysis, prepared the figures, and performed the statistical analysis. WR carried out the flow cytometry experiments. LS carried out the immunoassays and participated in the discussion. HB helped conceptualize the paper and participated in the discussions. DM performed the genetic screening. MA designed the study, coordinated the efforts, and drafted the manuscript. All authors read and approved the final manuscript.

## Acknowledgment

The authors want to acknowledge Ms. Xizi Hu for organizing the data and assisting with the flow cytometry experiments.

## Supplementary Information

*Supplement Figure 1. Flow cytometry histogram.* A flow cytometry histogram shows the IL-2 expression levels of the cells sorted in the final stage of the Jurkat T cells CRISPR screen.

*Supplement Figure 2. PTPN2 is required for PD-1 functions.* The percent of inhibition of secreted IL-2 from shirt hairpin (sh) scramble, and sh PTPN2 Jurkat T cells occulted overnight with Raji B cell vs. Raji B cell that overexpressed PD-L1 at different concentrations of SEE (A). p<0.05, n=5. ELISA measured IL-2 levels. Flow cytometry quantifies CD69 expression inhibition from sh Scramble and sh PTPN2 Jurkat T cells occulted overnight with Raji B cell vs. Raji B cell that overexpressed PD-L1 in the presence of SEE (B). Two different shRPTN2 clones are shown at different concentrations of SEE (A). p<0.05, n=5.

*Supplement Figure 3. FUT6 signaling downstream of the T cell receptor.* Growth assay of the different clones of short hairpin stably expressing Jurkat T cells stimulated with anti-CD3 and anti-CD28 over three days (A). Quantification of western blots showing the total ERK and phosphorylated ERK levels in the indicated sh Jurkat T cell lines stimulated with beads coated with antiCD3 for the appropriate time, as indicated in Figure (B). Fold change in secreted IL-2 levels from anti-CD3 stimulated shirt hairpin different Jurkat T cell lines measured by ELISA after overnight stimulation (C). n=3-5, * p<0.05. Cartoon showing the glycosylation pattern on FUT6 protein (D). Flow cytometry quantification of cell surface CD69 from Jurkat T cell line stably expressing the indicated sh and stimulated as indicated in Figure (E).

*Supplement Figure 4. The genes used to validate the dysfunctional T cell scores.* Heatmaps showing differential expression levels of genes that we discovered to be involved in PD-1 downstream signaling and that are also differentially expressed in the other scRNA sequencing datasets (GSE120575, SCP1288, and GSE164237), as reported in Fig. 2D. The green columns represent non-responders. The brown columns represent responders (A). The top gene list in each square is shown on the right side of the heat map. Volcano plot based on data from scRNA sequencing datasets (GSE120575, SCP1288, and GSE164237) displaying fold change in the expression levels of genes that were either up or down-regulated in the cancer patients that failed to respond to PD-1 blockade (B).

*Supplement Figure 5. Top-ranked genes are downregulated in T cells of other autoimmune diseases.* Expression levels on top-ranked genes that we discovered as needed for PD-1 signaling were upregulated in cancer patients that responded to PD-1 inhibition in other autoimmune T cell bult RNA sequencing (A). Both the T cell and non-T cell datasets were analyzed. Heatmap showing genes that have high T cell dysfunctional score (upper right box) and their expression levels in ICI datasets, immunosuppress datasets, and in our CRISPR screen (lower right box) (B). The list of top genes in each square is shown on the right side of the heat map. Representative genes plotted as box plots showing the mean RNA expression levels in SF RA vs. PB RA (Ci) or PB RA vs. PB HC (Cii).

*Supplement Figure 6. Enrichment analysis of transcription factors.* Heatmaps showing the predicted expression levels of transcription factors (TF) in different activated T cell subsets (CD4^+^ PRF1^+^, CD8^+^ HLA-DR^HIGH^, and CD8^+^ HLA-DR^LOW^ T cells) based on top-ranked genes that we discovered in the screen to be involved in PD-1 signaling and the context of PD-1, LAG3, and CTLA-4 (shown in purple) (A). Genes we discovered to be involved in PD-1 signaling are shown in pink. Expression levels in both PB and SF are shown. TF expression in PB, SF, and other T cell subsets (B). Spider plot depicting the enrichment of TF that are associated with the genes of CD8^+^ PD-1^+^ KLRG1^LOW^ T cells in SF (B).

