## Supplementary Figures for "PD-1 signaling uncovers a pathogenic subset of T cells in inflammatory arthritis"

#### Supplement Figure 1

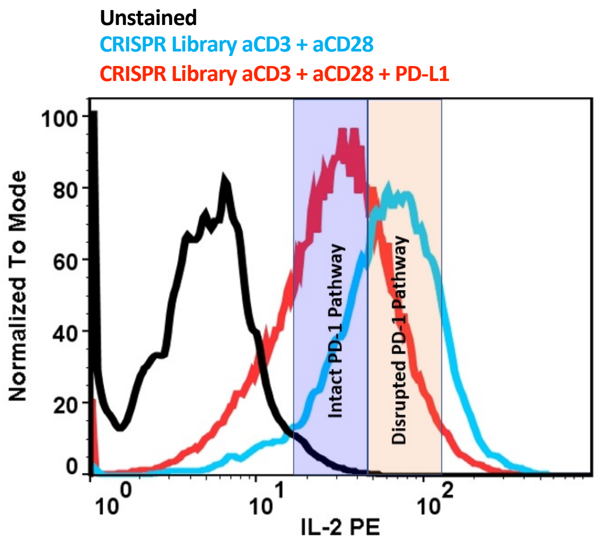

### Supplement Figure 2

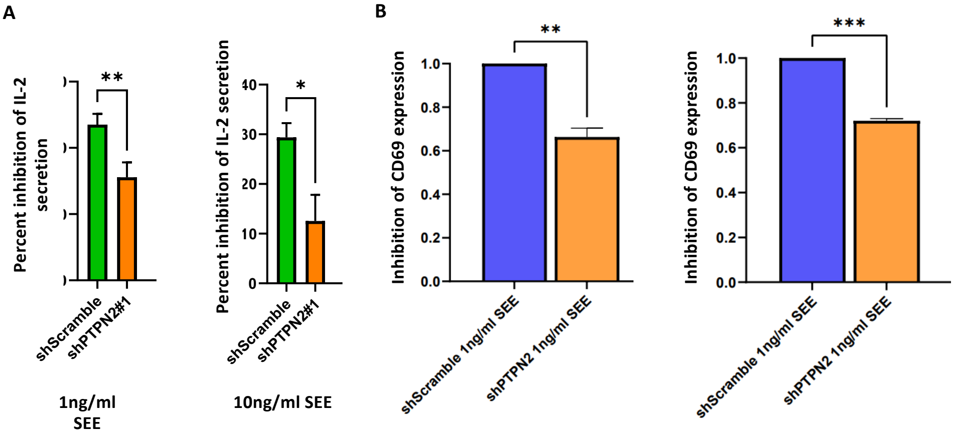

#### Supplement Figure 3

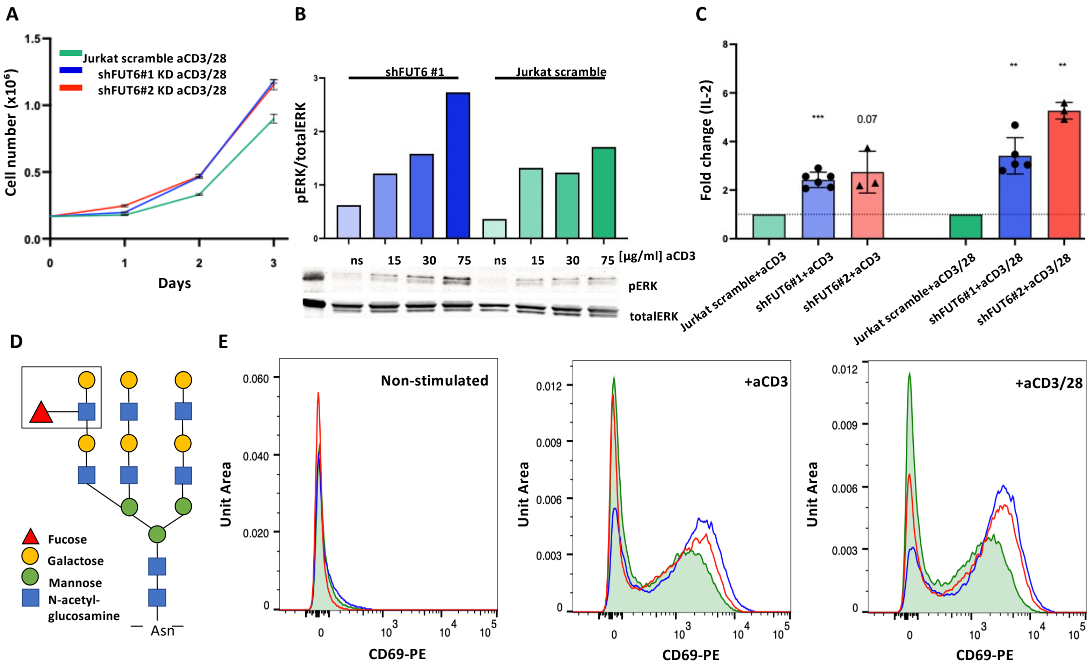

### Supplement Figure 4

A

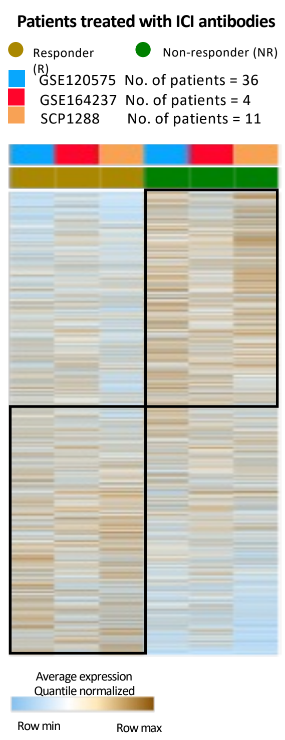


| | | | | | | | |
|---|---|---|---|---|---|---|---|
| FTH1P18 | NTS | SCGB1D2 | CXCL6 | PTH1R | KRT27 | C20ORF78 | NPR3 |
| RGS5 | KIR2DL3 | PDZK1 | BAIAP2L2 | RAD54L | CRYGC | APOL4 | SLCO4C1 |
| C17ORF97 | CST6 | GSTA2 | TNFRSF8 | OLFML3 | PTN | HMMR | VWA7 |
| PHACTR1 | RNF43 | SSTR1 | CDC45 | GPX6 | KCND3 | TRIO | PFN3 |
| KIF2C | PRLH | KRTAP6-2 | NCF2 | AP5B1 | SPACA3 | SULT2B1 | ODAM |
| LYZL1 | TEX11 | ALDH8A1 | TRAM1L1 | BASP1 | RHD | MEIS3 | GRXCR2 |
| ASB14 | ANLN | TSPAN1 | TMEM217 | CATSPER4 | ROR1 | GMPR | ZNF805 |
| SPDYE5 | GLIPR1L1 | GDA | BSX | CSF2RB | PNMA6A | DNMT3L | SDK1 |
| RIBC1 | SMPDL3B | SUN5 | KIR2DL1 | KCNJ14 | KANK4 | ZNF667 | BPIFB3 |

| | | | | | | | | |
|---|---|---|---|---|---|---|---|---|
| PABPC5 | SYT6 | ZNF492 | RND3 | LYG1 | PCDHGA1 | SMAD9 | SPRED3 | STC2 |
| SEMA5B | OR7G2 | CTRL | CADM2 | RAVER2 | VCAN | OR6K2 | CDH5 | FCGR1A |
| JAK2 | APC2 | CSF2RA | C2ORF80 | EVC2 | FAM71A | NEUROD1 | MTMR11 | ACTA1 |
| GJC1 | CACNA2D2 | TMEM47 | PSD2 | SLC9A4 | GPR176 | GPR179 | NANOS3 | PPP1R14C |
| IRX2 | ACKR3 | ASGR1 | CARD9 | TACR3 | C17ORF78 | CELF4 | APCS | HORMAD1 |
| TRPV6 | NRCAM | FBN3 | FST | TRPC5 | CCKBR | FLT1 | CXORF27 | FAM229A |
| GPX2 | GYPA | ZDHHC15 | KCNE3 | PNMT | HOXB5 | NOXRED1 | MAGEA2 | OR10AD1 |
| OR7D2 | FNDC4 | GSTT2 | CLCN4 | LTBP2 | PLCD4 | OVOL2 | RIPK4 | CDK18 |
| DLC1 | H1FNT | NOTCH3 | SHISA9 | PODN | TCL1B | KCTD1 | FUT6 | CHRNG |
| TCF21 | P2RY1 | C5ORF47 | ABHD12B | MEOX2 | S100A3 | ACTL9 | LAMP5 | RBM46 |
| CLLU1OS | CD300LG | UNC5C | PPFIA4 | STMND1 | LCE5A | KCNJ5 | MIOX | FAM107A |
| CAMK2B | AP3B2 | RFX4 | HOXD1 | | | | | |

B

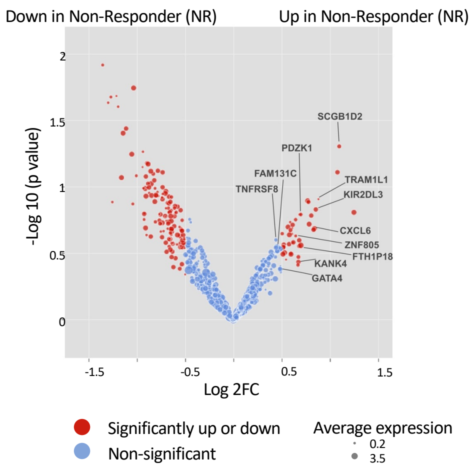

### Supplement Figure 5

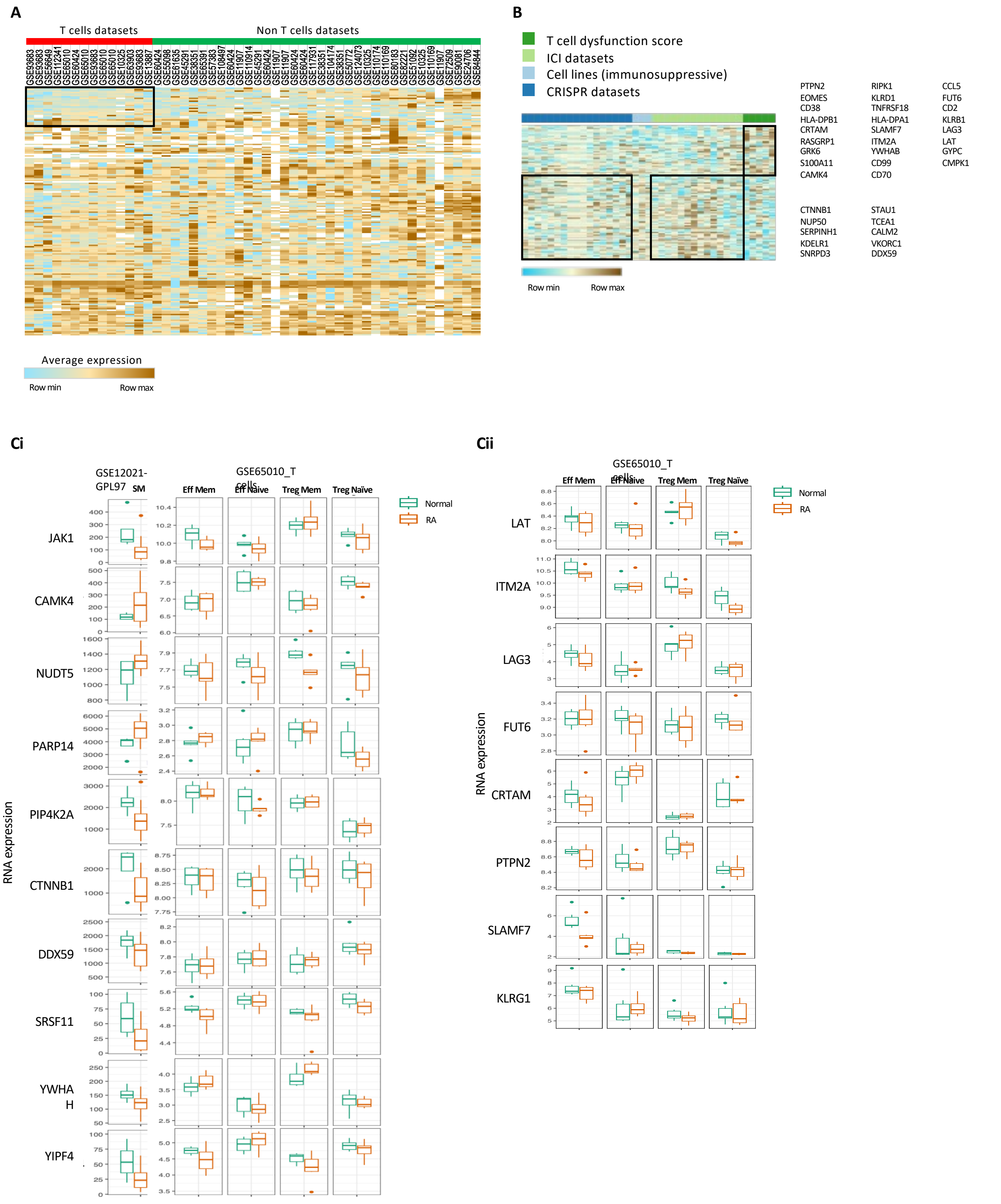

#### Supplement Figure 6

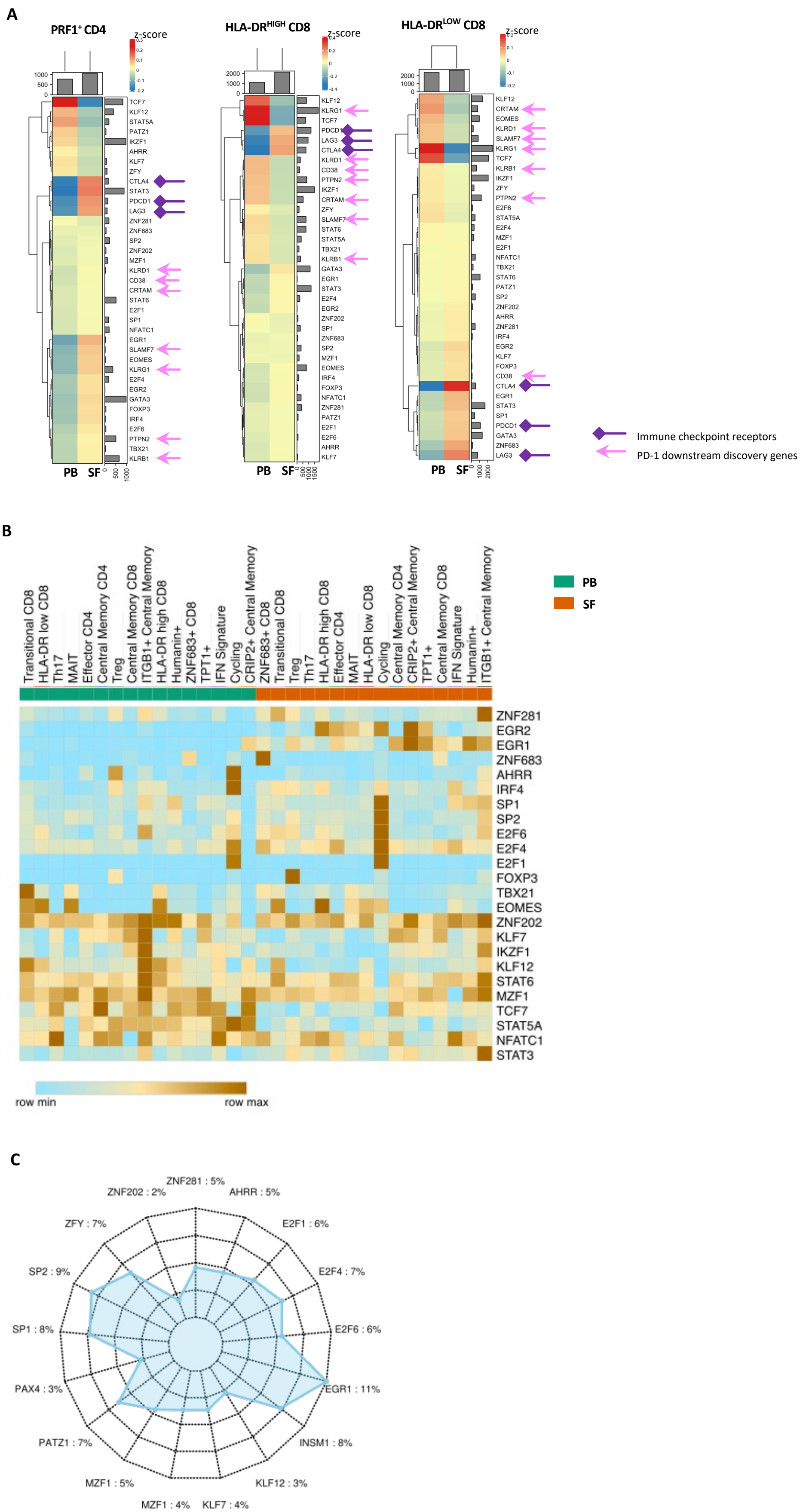
